# Resource-efficient Neural Network Architectures for Classifying Nerve Cuff Recordings on Implantable Devices

**DOI:** 10.1101/2022.10.05.510983

**Authors:** Y. E. Hwang, R. Genov, J. Zariffa

**Author notes:** **Disclosures: Y. E. Hwang:** None. **R. Genov:** None. **J. Zariffa:** None.

## Abstract

**Background:** Closed-loop control of functional electrical stimulation involves using recorded nerve signals to make decisions regarding nerve stimulation in real-time. Surgically implanted devices that can implement this strategy have significant potential to restore natural movement after paralysis. Previous work demonstrated the use of convolutional neural networks (CNNs) to discriminate between activity from different neural pathways recorded by a high-density multi-contact nerve cuff electrode. Despite state-of-the-art performance, that approach required too much data storage, power and computation time for a practical implementation on surgically implanted hardware.

**Objective:** To reduce resource utilization for an implantable implementation, with a minimal performance loss for CNNs that can discriminate between neural pathways in multi-contact nerve cuff electrode recordings.

**Methods:** Neural network (NN) architectures were evaluated on a dataset of rat sciatic nerve recordings previously collected using 56-channel (7 x 8) spiral nerve cuff electrodes to capture spatiotemporal neural activity patterns. The NNs were trained to classify individual, natural compound action potentials (nCAPs) elicited by sensory stimuli. Three architecture types were explored: the previously reported ESCAPE-NET, a fully convolutional network, and a recurrent neural network. Variations of each architecture yielded NNs with a range in the number of weights and required floating-point operations (FLOPs). Each NN was evaluated based on F1-score and resource requirements.

**Results:** NNs were identified that, when compared to ESCAPE-NET, required 1,132-1,787x fewer weights, 389-995x less memory, and 6-11,073x fewer FLOPs, while maintaining macro F1-scores of 0.70-0.71 compared to a baseline of 0.75. Memory requirements range from 22.69 KB to 58.11 KB, falling within the range of on-chip memory sizes from several published deep learning accelerators fabricated in 65nm ASIC technology.

**Conclusion:** Reduced versions of ESCAPE-NET require significantly fewer resources without significant accuracy loss, thus can be more easily incorporated into a surgically implantable device that performs closed-loop real-time responsive neural stimulation.

## 1 Introduction

Afferent signals contain proprioceptive and sensory information, which can be used to determine joint position, muscle stretch and tension, and presence of external stimuli. The central nervous system (CNS) uses such information to guide limb movement and reflex responses [1]; in the case of paralysis due to spinal cord injury, supraspinal control of limb movements may be disrupted, preventing the muscles from receiving motor commands from the brain.

Open-loop functional electrical stimulation (FES) can be delivered via peripheral nerves to restore motor function in paralyzed limbs, but without the sophistication of motor commands generated by the brain. This results in coarse limb movement lacking the nuance of unimpaired motor control, the inability to compensate for unexpected perturbations or errors, and inefficient stimulation delivery that causes muscle fatigue [2]-[4]. Closed-loop FES aims to improve stimulation delivery by using afferent signals as feedback to regulate the stimulation delivered.

The information-carrying afferent signals can be recorded using nerve interfacing electrodes, after which data processing can be applied to extract and interpret the proprioceptive and sensory information. Several studies have demonstrated using such information as feedback control to achieve closed-loop FES in animal models. Bruns et al. [5] demonstrated joint angle decoding by applying linear regression on dorsal root ganglion (DRG) signals to regulate FES of various leg muscles to produce a regular gait cycle. Holinski et al. [6] demonstrated ground reaction force prediction during stepping by multivariate linear equations on DRG signals to regulate intraspinal microstimulation to control hind leg motion. Song et al. [7] demonstrated ankle angle estimation by fast independent component analysis and dynamically driven recurrent neural network on sciatic nerve signals to regulate FES to control ankle motion. Closed-loop FES has also been demonstrated in human trials. Wendelken et al. [8] demonstrated closed-loop control of a virtual prosthetic hand using intraneural electrodes on the human median nerve decoded by a Kalman filter. Hansen et al. [9] demonstrated closed-loop foot drop correction using human sural nerve recordings discriminated by an artificial neural network. Inmann et al. [10] demonstrated a closed-loop hand grasp system using human median nerve recordings processed by rectification and bin-integration, high- and low-pass filtering, and delay to control hand grasp to react to and stop slipping objects.

Peripheral nerve signals may be recorded by surgically implanted devices of varying levels of invasiveness [11]. Extraneural nerve cuffs are advantageous as they do not penetrate the nerve and are typically preferable for clinical use due to long-term stability [12], [13]. However, naturally-evoked compound action potentials (nCAP) exhibit low signal-to-noise ratio (SNR). Thus, achieving recording selectivity (i.e. the ability to determine which neural pathway produced the measured signals) using extraneural devices is challenging [11]. Improving the selectivity of peripheral nerve recordings is necessary to provide functionally specific feedback signals that can improve the performance of closed-loop FES systems.

Koh et al. recently demonstrated that selective recording of individual nCAPs from extraneural nerve cuff electrodes was possible. Deep learning was used to classify sciatic nerve recordings obtained by multi-contact nerve cuffs with state-of-the-art selectivity [14]. They used multi-contact nerve cuffs with multiple rings of contacts along the length of the nerve and multiple contacts per ring around the diameter of the nerve; this structure captures velocity and spatial variability, respectively, between different neural pathways. The novel method, named Extraneural Spatiotemporal CAPs Extraction Network (ESCAPE-NET), detects individual nCAPs then represents them as a 2-dimensional spatiotemporal signature. ESCAPE-NET uses a convolutional neural network (CNN) to classify spatiotemporal signatures. Classifying individual nCAPs is particularly advantageous as it provides high temporal resolution and could potentially handle scenarios where multiple neural pathways are active, which complicates classification when using longer time windows.

To be successfully translated to humans, closed-loop FES technology must be implemented on surgically implantable devices. A surgically implantable device would exhibit minimal interference with a user’s daily routine and would draw minimal attention to the person’s disability due to being externally invisible [15] [16]. However, neural networks (NN) traditionally rely on computational complexity to achieve high classification accuracy for non-linearly separable datasets. Computational complexity translates to longer computation time and size of hardware in terms of either memory, accelerators, or power consumption, which hinders deployment onto surgically implantable devices with strict hardware constraints. Recent efforts on reducing a NN’s resource intensiveness with minimal performance degradation include MobileNet [17] [18] [19], EfficientNet [20] [21], and Once-For-All [22] networks. Specifically, focus has been placed on reducing the number of parameters or weights, which are crucial to the NN’s capacity to learn and detect structural features in the input data. Springenberg et al. developed a fully convolutional network that outperformed more complicated state-of-the-art networks, achieving large parameter savings that doubles as a form of regularization [23]. Another benefit of reducing the computational burden of a NN is the reduction of latency (computational time for inference) required for a NN to process its inputs.

To combine the proven benefits of using NNs to discriminate between neural signals and the goal of real-time performance on implantable hardware, there is a need to explore reduced NNs that perform with minimal accuracy loss and are suitable for real-time performance on implantable hardware. This study aimed to reduce resource consumption with minimal performance accuracy loss for machine learning techniques that can distinguish between neural pathways in multi-contact nerve cuff electrode recordings.

## 2 Methods

### 2.1 Dataset description

The comparison of the neural network architectures made use of an existing dataset consisting of spatiotemporal signatures extracted from detected nCAPs in the rat sciatic nerve. More details regarding the dataset and how it was collected can be found in [24]. Figure 1 illustrates detected nCAPs and their corresponding spatiotemporal signatures. There are three classes of spatiotemporal signatures in this dataset, labeled based on the type of stimulus that was being applied at the time each nCAP was recorded: ankle dorsiflexion, ankle plantarflexion, and cutaneous stimulus to the heel using a Von Frey monofilament. Data was collected from 9 Long-Evan rats, with a separate neural network trained for each rat. 3-fold cross-validation was used in each case.

**Figure 1.**
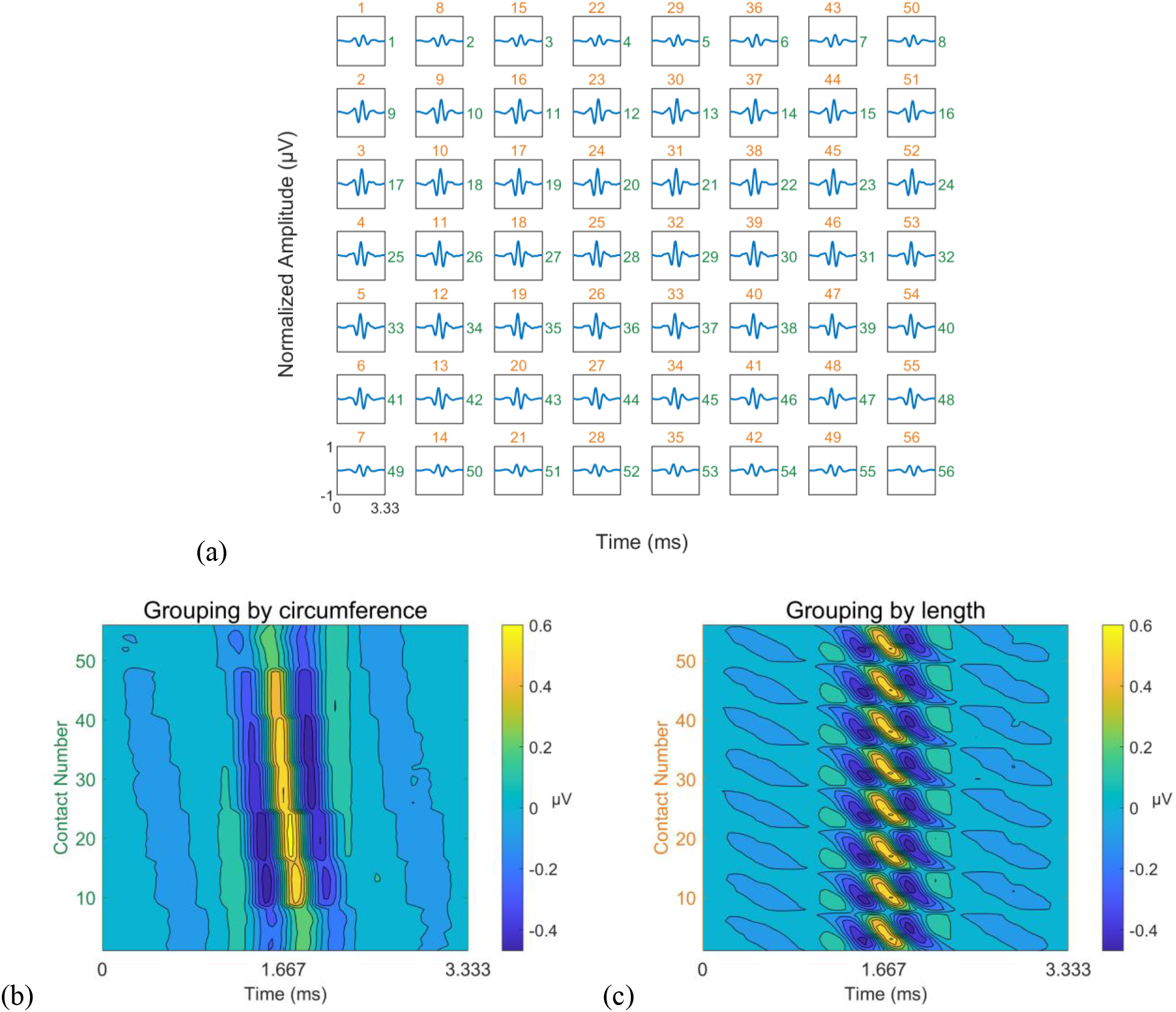
Example of the ENG signals used for classification. For clarity of visualization, the depicted signals are averaged across several thousands of nCAPs. Inputs to the NNs consisted of single nCAP recordings, which would contain higher levels of noise. (a) Processed signals of a detected nCAP measured on all contacts of the nerve cuff electrode. The signals have been filtered, tripole referenced and normalized between −1 and 1 before being averaged. (b) Spatiotemporal signature representation of the ENG of the signals presented in (a); contacts are numbered based on the green numbers shown in (a) to the right of each plotted nCAP. (c) Spatiotemporal signature representation of the ENG of the signals presented in (a); contacts are numbered based on the orange numbers in (a) on top of each plotted nCAP.

Electroneurographic (ENG) signals were recorded at 30 kHz using the 56 contacts of the nerve cuff electrode, which were arranged as 7 rings of 8 contacts. Each spatiotemporal signature contained a 3.33 ms segment of the recorded signal, centered around the detected peak in the middle ring of contacts. Each spatiotemporal signature therefore had dimensions of 56 channels x 100 time samples. Two representations of the spatiotemporal signature are possible: the first groups channels by nerve circumference (i.e. every 8 consecutive rows correspond to a ring of contacts) and the second groups channels by nerve length (i.e. every 7 consecutive rows correspond to a line of contacts along the nerve).

The number of nCAPs per category was 3,898 ± 2,175 for dorsiflexion, 3,454 ± 2,433 for plantarflexion, and 3,991 ± 2,145 for heel prick. Training sets for each rat were augmented to include a total of 10,000 training examples of spatiotemporal signatures per class. Augmented nCAPs were generated by averaging all nCAPs associated with a label then adding randomly sampled noise to create new noisy nCAPs [14]. Testing sets included only detected nCAPs (no augmented nCAPs). The validation set was comprised of a randomly sampled 10% of the training set.

### 2.2 Evaluation of neural networks

Several NN architectures were explored for the purpose of classifying spatiotemporal signatures and evaluated based on the amount of required resources and classification ability. The goal was to determine whether or not there exists a NN architecture that uses an amount of resources compatible with eventual application-specific integrated circuit (ASIC) translation while demonstrating competitive classification performance.

For the purposes of this study, we define resources as a network’s required memory and floating-point operations (FLOPs), which are directly influenced by the network’s architecture, the number of weights in a network, and the precision of the numeric representation of the weights and intermediate values. We define a network’s classification ability based on accuracy and macro F1-score of the network’s classification of a labelled test set. For each datum in the test set, the label with the highest softmax output of the neural network is the predicted label.

Number of parameters was measured using the built-in Keras function *count_params*. Amount of memory was measured using the size of Tensorflow Lite (TFLite) converted models. FLOPs were measured using the Tensorflow Profiler. When determining classification accuracy and F1-score, the label with the highest predicted probability is compared to the ground truth label for equivalence.

### 2.3 Implementation and training of neural networks

All neural network models were designed and trained using Tensorflow Keras libraries. Training was performed over 1000 epochs or whenever the validation loss function did not decrease for over 15 epochs. The optimization algorithm used for network training was stochastic gradient descent (SGD) with a momentum value of 0.9. The loss function used was categorical cross entropy (CCE) loss. All networks ended with a dense layer responsible for classification using the softmax activation function.

### 2.4 Neural network architectures

We refer to an architecture as a specific type of network. The first architecture was ESCAPE-NET, the CNN previously published by Koh et al. [14]. The second architecture was a Fully Convolutional Network (FCN), as inspired by Lin et al. [25], using smaller filter sizes and deeper networks as inspired by multiple recent successful neural networks [26]. Fully convolutional networks replace traditional dense layers with convolutional layers that perform the same functionality and have been shown to reduce overfitting and promote parameter efficiency. The third architecture was a Recurrent Neural Network (RNN), which was explored to exploit the time series nature of the dataset in question.

Within a single architecture, structural details were varied to explore networks with different trade-offs between amount of resources and performance. Examples include the network width (i.e. the number of filters per layer), the number of layers, filter sizes, regularization, or types of layers within the same family (e.g. Long-Short Term Memory and Gated Recurrent Unit layers for the RNN family).

Each structural detail was explored sequentially. After each structural detail was explored, the network that yielded the fewest number of weights while maintaining performance higher than 0.7 macro F1-score was chosen. Each structural detail remained fixed based on this requirement for the remainder of the exploration.

#### 2.4.1 ESCAPE-NET Mini

Introduced by Koh et al. [14], ESCAPE-NET receives two inputs, corresponding to two different representations of the spatiotemporal signature. The two representations differ in the ordering of the contacts. The “spatial emphasis” representation ensures that rows of pixels are locally related based on contact placement around the diameter of the nerve, whereas the “temporal emphasis” representation ensures that rows of pixels are locally related based on contact placement along the length of the nerve (Figure 1). The former exploits the spatial variation of different neural pathways, and the latter exploits the velocity variation of different neural pathways. Reduced versions of ESCAPE-NET, hereon noted as ESCAPE-NET Mini, are explored by varying the following structural details detailed in this section.

For each of the two inputs of the network, there are multiple convolutional layers using the ReLU activation function in sequence, separated by 2×2 max pooling layers for data downsampling. To initialize the weights of the first and second convolutional layers, they were temporarily connected to dense layers and trained for 25 epochs. The pair of convolutional layer stacks were concatenated after the third convolutional layer into a dense layer using the ReLU activation function. Figure 2a illustrates a sample ESCAPE-NET network explored in this work.

**Figure 2.**
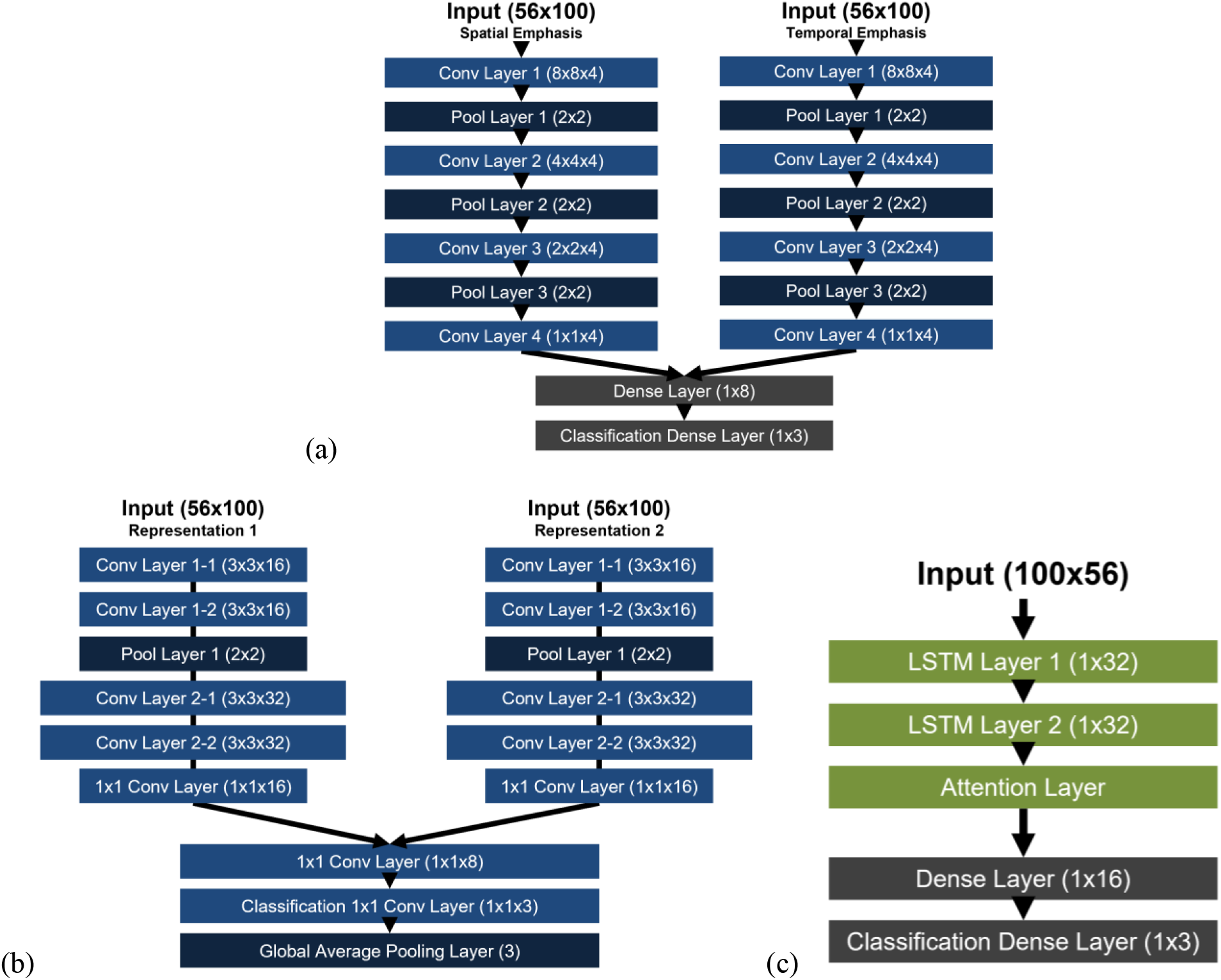
Network design for the novel architectures explored: (a) ESCAPE-NET Mini architecture, (b) ESCAPE-FCN architecture, and (c) ESCAPE-RNN architecture. Structural detail values used in these illustrations are based on the highlighted networks presented in the Results section.

An initial sweep of network widths was performed by varying the convolutional layer width and dense layer width between 2-64, increasing by a factor of 2, keeping the two values equal throughout this sweep. A specific value was selected after this initial sweep using the network with the fewest weights performing above a macro F1-score of 0.7. During this sweep, the number of convolutional layers was set to 3, the filter sizes were set to 8×8 for conv layer 1, 4×4 for conv layer 2, 2×2 for conv layer 3, and the dropout value was set at 0.

To investigate the effect of unequal convolutional and dense layer widths, a secondary sweep of network widths was performed on a smaller search space centered around the chosen value from the initial sweep. For example, if a width of 8 was chosen after the initial sweep, convolutional and dense widths were explored for all combinations of 4, 8 and 16 (i.e. 9 combinations in total). Specific values were selected for convolutional and dense layer widths after this secondary sweep.

The number of layers was varied between 2-4. Filter sizes were varied between 8×8 for conv layer 1, 4×4 for conv layer 2, 2×2 for conv layer 3 and 9×9 for conv layer 1, 5×5 for conv layer 2, 3×3 for conv layer 3. The dropout value was varied between 0 and 0.5 to represent the absence and presence of a dropout layer, respectively. The dropout value of 0.5 was used due to its reported efficacy [27].

#### 2.4.2 Fully Convolutional Network (FCN)

The second explored architecture was an FCN using smaller filter sizes and a deeper network. Like ESCAPE-NET, it receives two inputs for the different representations of the spatiotemporal signature. For each of the two inputs of the network, there are multiple sets of convolutional layers using the ReLU activation function and filter size of 3×3 in sequence, separated by 2×2 max pooling layers for data downsampling. The second and third sets of convolutional layers were followed by a 1×1 convolutional layer to allow for more complex function modelling. The pair of convolutional layer stacks were concatenated after the third set of convolutional layers into a 1×1 convolutional layer using the ReLU activation function in place of the dense layer in ESCAPE-NET, and a 1×1 convolutional layer and Global Average Pooling layer in place of the dense layer responsible for classification. Figure 2b illustrates a sample FCN network explored in this work.

An initial sweep of network widths was performed by varying the number of filters of the first set of convolutional layers (i.e. base convolutional layer width) between 2-64, increasing by a factor of 2, and varying the number of filters of the 1×1 layer after concatenation (i.e. 1×1 convolutional layer width) between 2-64, increasing by a factor of 2, keeping the two values equal throughout this sweep. A specific value was selected for each width after these initial sweeps based on its resource/performance trade-off. During this initial sweep, the number of sets of layers was set to 3, and the channel width multiplier was set to 2. The channel width multiplier determines how layer width increases from one set of convolutional layers to the next.

To investigate the effect of unequal base convolutional layer and 1×1 convolutional layer widths, all combinations of the selected width, half the selected width and double the selected width were evaluated (i.e. 9 networks in total). A specific value was selected for each width after this exploration.

Structural details that were varied for this architecture were the number of sets of layers and channel width multiplier. The choice to vary these particular structural details was based on CNN advancements that found success using deeper networks with smaller filter sizes [28], [29], increasing width of layers in deeper layers [26], and 1×1 convolutions in the place of dense layers [25].

The number of sets of convolutional layers was varied between 2-4. The channel width multiplier value was varied between 1-3.

#### 2.4.3 Recurrent Neural Network (RNN)

The third explored architecture was an RNN consisting of multiple recurrent layers using the tanh activation function connected in sequence, followed by a dense layer using the ReLU activation function. Unlike the previous architectures, there is only one input to this network, as each contact of the nerve cuff is treated as a single channel of the data, eliminating the benefit of having localized spatial or temporal information encoded in the spatiotemporal signature. Figure 2c illustrates a sample RNN network explored in this work.

An initial sweep of network widths was performed by varying the number of units of the recurrent layer (i.e. recurrent layer width) and the dense layer width between 2-64, increasing by a factor of 2, keeping the two values equal throughout this sweep. A specific value was selected for each width after these initial sweeps based on its resource/performance trade-off. During these sweeps, the number of layers was set as 3, and the layer type was set as GRU.

To investigate the effect of unequal recurrent layer and dense layer widths, all combinations of the selected width, half the selected width and double the selected width were evaluated (i.e. 9 networks in total). A specific value was selected for each width after this exploration.

Structural details that were varied for this architecture were the number of layers, number of neurons per layer, and layer type. The number of layers of the network was varied from 2-5 given a fixed number of neurons per layer. The number of layers was fixed after this exploration. Layer types that were explored consisted of Long-Short Term Memory (LSTM), Gated Recurrent Unit (GRU), Bidirectional LSTM, and Bidirectional GRU. The last layer was varied between the recurrent layer and an attention layer.

### 2.5 Lower weight precision and memory requirement definition

Lower precision numerical representation of weights was explored due to its direct relevance to minimizing hardware requirements. Using fewer bits to represent a numerical value reduces the amount of memory required to store values and the complexity of multiply and accumulate operations, but potentially sacrifices accuracy due to information loss. By default, Tensorflow models use 32-bit floating point numbers to store weights. We explored using 8-bit integer weights using the TensorFlow Lite default optimization from the TensorFlow Model Optimization Toolkit. A TFLITE file is produced that contains the neural network model’s execution graph. We used the size of a TFLITE file to define a network’s memory requirement.

## 3 Results

We present the results of the network exploration performed by varying structural detail values for each of the three architectures. Each network’s classification accuracy, macro F1-score, number of weights and FLOPs are presented. For each architecture, a single network is highlighted and analyzed in more detail.

Figure 3 displays the calculated number of weights for all the explored networks for each of the three architectures measured against their achieved macro F1-score. Figure 4 displays the calculated number of FLOPs for the same networks against their achieved macro F1-score. A single datapoint represents the average performance of a network across all 9 subjects and 3 folds. A highlighted network, as indicated by the markers with black borders, was selected for each of the three architectures based on the architecture requiring the fewest weights that meets our performance requirement of 0.70 macro F1-score.

**Figure 3.**
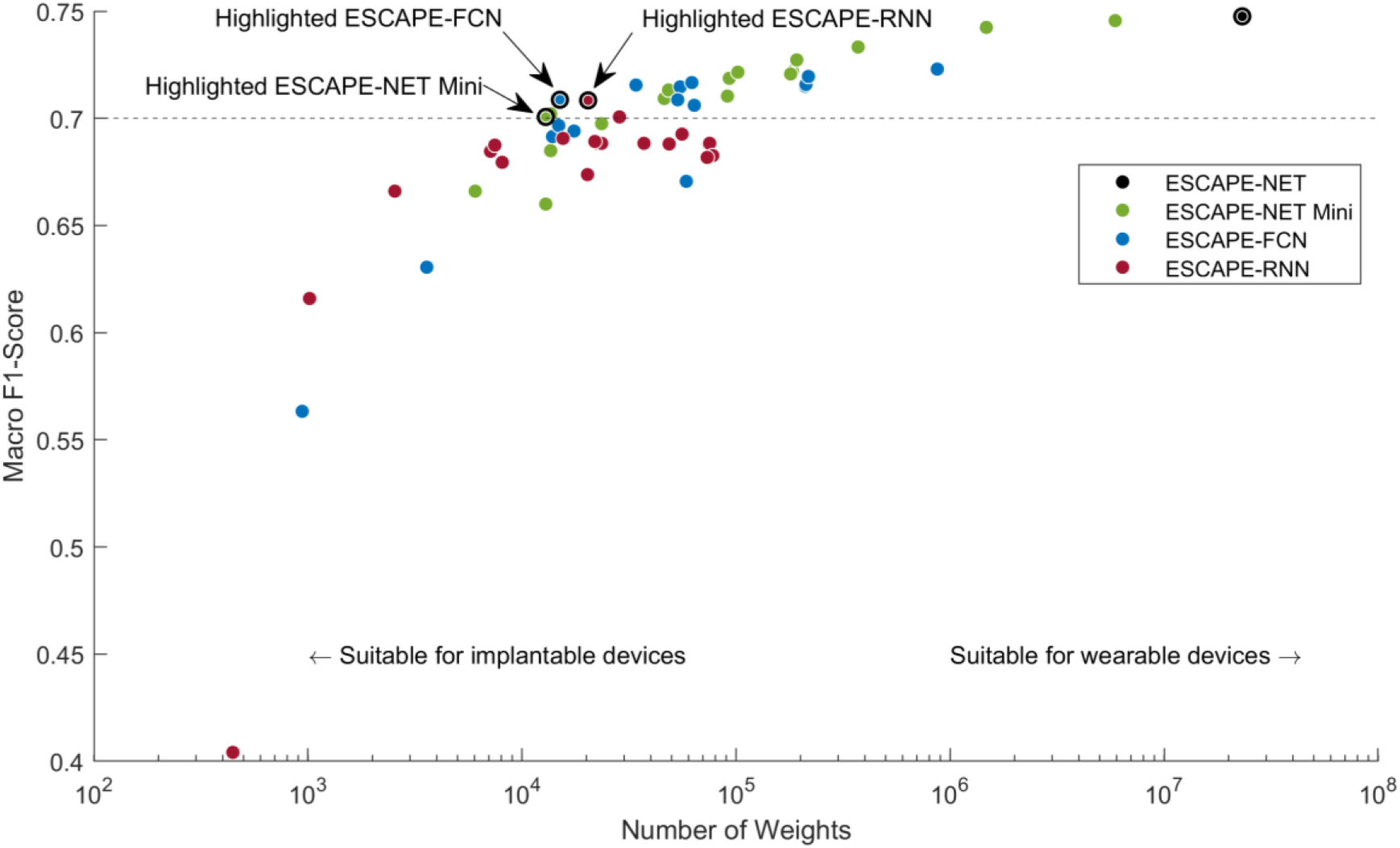
Resource/performance trade-off plot for every explored network configuration for each of the three architectures, analyzing the macro F1-score for performance versus the number of weights for resources. Each data point represents the average performance of a network design trained with 27 datasets (9 subjects, 3-fold cross validation). The dotted line represents the minimum performance constraint used in this study. The points with black borders represent the highlighted network configurations selected for each of the 3 architectures as well as the original ESCAPE-NET architecture.

**Figure 4.**
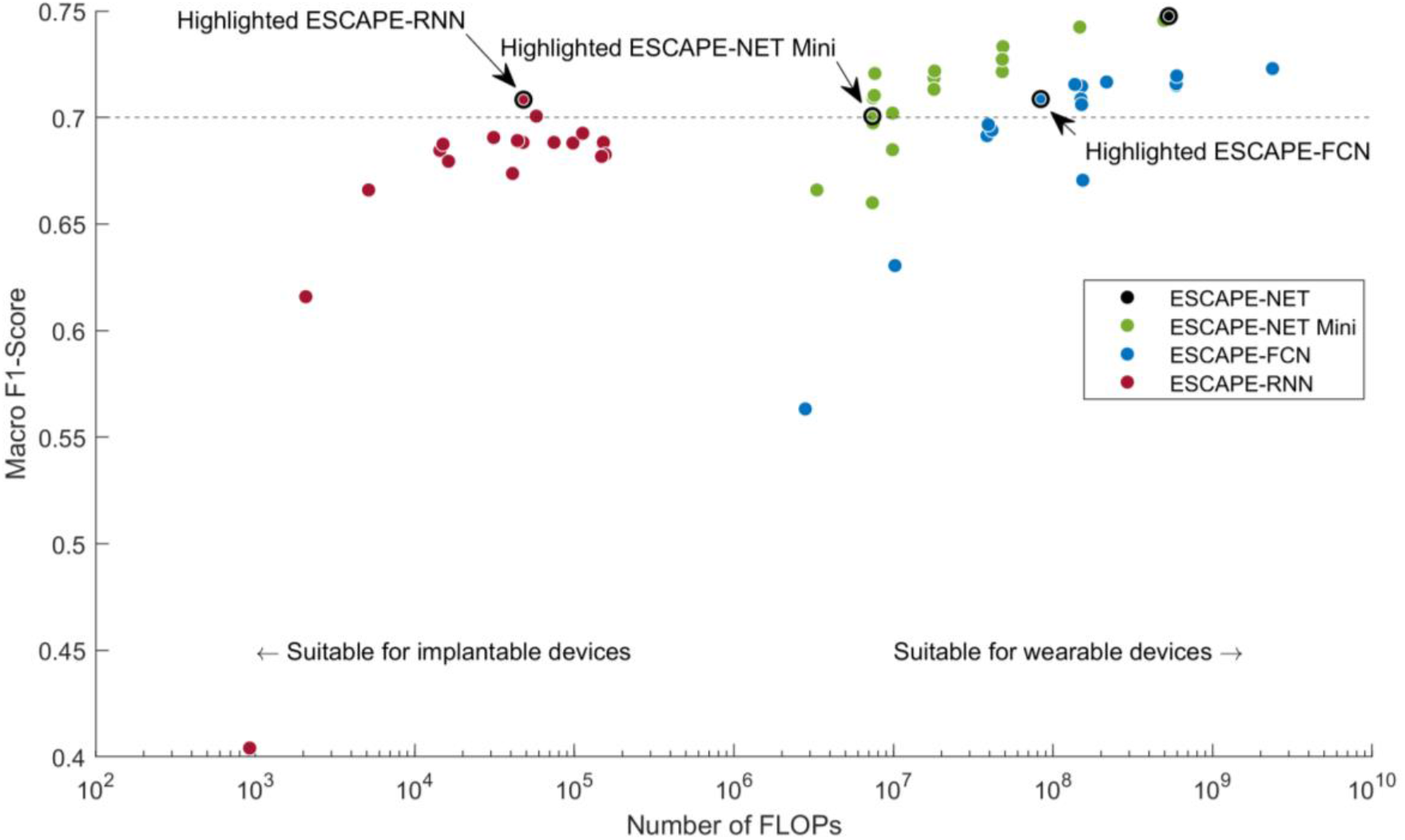
Resource/performance trade-off plot for every explored network configuration for each of the 3 architectures, analyzing the macro F1-score for performance versus the number of FLOPs for resources.

As expected, the performance of the networks within each architecture trended downwards as the number of resources decreased. The majority of the explored networks performed within 0.10 macro F1-score of the baseline network from [14].

### 3.1 ESCAPE-NET Mini architecture

The initial sweep of layer widths was performed by varying the convolutional and dense layer width values between 2, 4, 8, 16, 32, and 64, keeping the two values equal. The convolutional and dense layer widths were chosen to be 8 after this initial sweep. The secondary sweep of layer widths was performed by varying the convolutional and dense layer width values between 4, 8 and 16, exploring all combinations. The convolutional layer width was chosen to be 4 and the dense layer width was chosen to be 8 after this secondary sweep.

The number of convolutional layers was chosen to be 4 after exploring networks with 2 layers, 3 layers and 4 layers. The first layer uses 8×8 filters, the second layer uses 4×4 filters, the third layer uses 2×2 filters, and the fourth layer uses 1×1 filters. Dropout was not used after the dense layer.

### 3.2 ESCAPE-FCN architecture

The initial sweep of layer widths was performed by varying the base convolutional and 1×1 convolutional layer width values between 2, 4, 8, 16, 32, and 64, keeping the two values equal. The base convolutional and 1×1 convolutional layer widths were chosen to be 16 after this initial sweep. The secondary sweep of layer widths was performed by varying the base convolutional and 1×1 convolutional layer width values between 8, 16 and 32, exploring all combinations. The base convolutional layer width was chosen to be 16 and the 1×1 convolutional layer width was chosen to be 8 after this secondary sweep.

The number of sets of convolutional layers was chosen to be 2 after exploring networks with 2, 3 and 4 sets of layers. The channel width multiplier value was chosen to be 2.

### 3.3 ESCAPE-RNN architecture

The initial sweep of layer widths was performed by varying the recurrent and dense layer width values between 2, 4, 8, 16, 32, and 64, keeping the two values equal. The recurrent and dense layer widths were chosen to be 32 after this initial sweep. The secondary sweep of layer widths was performed by varying the recurrent and dense layer width values between 16, 32 and 64, exploring all combinations. The recurrent layer width was chosen to be 32 and the dense layer width was chosen to be 16 after this secondary sweep.

The number of layers was chosen to be 3. The 1^st^ and 2^nd^ layers were chosen to be LSTM, and the 3^rd^ layer was an attention layer.

### 3.4 Resource/Performance trade-off analysis

To analyze the resource/performance plots presented in the Results section, networks nearer the top left corner are preferable. However, this study is not strictly an optimization problem for minimal resources and maximum performance. Alternative ways of interpreting the plots are by defining a resource constraint given a specific hardware platform then selecting the network that achieves the best performance within this constraint or defining a minimum performance requirement and selecting the network that uses the fewest resources above this threshold.

To select a highlighted network for each architecture as presented in Figure 3, we use a performance constraint of a macro F1-score threshold of 0.7. In Figure 5b, we use resource constraints specified by published deep learning accelerator implementations. These resource and performance constraints are specific to this study. Different constraints may be defined based on different applications and platforms. Within this selection of networks, there are many alternatives to the highlighted networks that may be suitable for practical use.

### 3.5 Highlighted networks

Table 1 summarizes the evaluated metrics of the three highlighted networks. These are the networks for each architecture that require the fewest weights while performing with macro F1-score of above 0.70. Of the three selected networks, ESCAPE-NET Mini uses the fewest weights and therefore memory, and ESCAPE-RNN uses the fewest FLOPs. ESCAPE-NET Mini can achieve better performance at higher amounts of resources, whereas the other two architectures plateau.

**Table 1.** Summary of the performance and resource requirements of the baseline network and the 3 highlighted networks.

| Architecture | Classification Accuracy (%) | Macro F1-Score | Number of Weights | Memory (KB) | FLOPs |
| --- | --- | --- | --- | --- | --- |
| ESCAPE-NET [14] | **80.76 ± 10.13** | **0.75 ± 0.11** | 23,112,067 | 22,585 | 529,405,202 |
| ESCAPE-NET Mini | 77.22 ± 11.64 | 0.70 ± 0.13 | **12,931** | **22.69** | 7,405,290 |
| ESCAPE-FCN | 77.71 ± 10.79 | 0.71 ± 0.12 | 15,075 | 27.32 | 83,549,215 |
| ESCAPE-RNN | 78.41 ± 10.01 | 0.71 ± 0.12 | 20,423 | 58.11 | **47,810** |

Table 2 compares the performance and memory for the three highlighted networks with and without lower precision weights. Every network performs with approximately equal accuracy and macro F1-score after conversion to lower precision weights. All reported instances of memory footprint in this document outside of Table 2 uses the TF optimized model with 8-bit integer weights.

**Table 2.** Performance and memory comparison between higher- and lower-precision weights.

| Architecture | 32-bit floating point weights |  |  |
| --- | --- | --- | --- |
|  | Classification Accuracy (%) | Macro F1-Score | Memory (KB) |
| ESCAPE-NET [14] | $80.77 \pm 10.13$ | $0.75 \pm 0.11$ | 90,287 |
| ESCAPE-NET Mini | $77.22 \pm 11.64$ | $0.70 \pm 0.13$ | 57.65 |
| ESCAPE-FCN | $77.70 \pm 10.78$ | $0.71 \pm 0.12$ | 65.70 |
| ESCAPE-RNN | $78.41 \pm 10.01$ | $0.71 \pm 0.12$ | 113.05 |
|  | 8-bit integer weights |  |  |
|  | Classification Accuracy (%) | Macro F1-Score | Memory (KB) |
| ESCAPE-NET [14] | $80.76 \pm 10.13$ | $0.75 \pm 0.11$ | 22,585 |
| ESCAPE-NET Mini | $77.22 \pm 11.64$ | $0.70 \pm 0.13$ | 22.69 |
| ESCAPE-FCN | $77.71 \pm 10.79$ | $0.71 \pm 0.12$ | 27.32 |
| ESCAPE-RNN | $78.44 \pm 10.01$ | $0.71 \pm 0.12$ | 58.11 |

## 4 Discussion

This study explored 3 different NN architectures, varying structural details for each to explore the trade-off between performance and accuracy for networks with reduced resource requirements compared to the baseline network from [14]. We found versions of each architecture with significantly reduced resource requirements that maintained macro F1-score of above 0.70. In particular, the networks’ memory requirements fell by an order of magnitude from 10s of megabytes to 10s of kilobytes.

### 4.1 Comparison between highlighted networks

With structural detail tuning, a version of each architecture could achieve comparable trade-offs between the number of weights and macro F1-score. ESCAPE-NET Mini performs slightly better when given more weights, and ESCAPE-RNN performs slightly better when given fewer weights. These networks may be better choices if the hardware constraints in a practical setting are loosened or tightened, respectively. Hardware constraints may be loosened due to implementation of the neural network on a wearable device or cloud storage in communication with the implant.

However, when considering FLOPs, ESCAPE-RNN networks are clearly advantageous, and ESCAPE-FCN networks are clearly disadvantageous. If processing speed is a bottleneck, an ESCAPE- RNN network may be preferred.

### 4.2 Defining resource constraints

Whether a particular neural network can be implemented onto a particular target device with real-time performance is a complex question that is difficult to determine prior to actual implementation. However, as this work was intended to facilitate eventual hardware implementation, we are interested in defining an informed estimate of potential memory and latency constraints and analyzing the neural networks we presented that fall within these limitations. Based on specifications of published deep learning implementations on ASIC technology and the memory requirements of our neural networks, we argue that our neural networks can be implemented on 65nm technology.

Moons et al. achieved a low-power high-precision embedded processor designed for large scale CNNs implemented in 40nm technology and equipped with 148 KB of on-chip memory [32]. Both Qiao et al. and Yue et al. achieved Computing-in-Memory-based processors for NNs implemented in 65nm technology equipped with 73 KB [33] and 164 KB [31] of on-chip SRAM, respectively. Chen et al. achieved an energy-efficient accelerator for Deep CNNs implemented in 65nm technology and equipped with 181.5 KB of on-chip SRAM [30]. As illustrated in Figure 5b, each of this work’s proposed networks’ memory requirements fall within the amount of on-chip SRAM available on these neural network hardware implementations.

**Figure 5.**
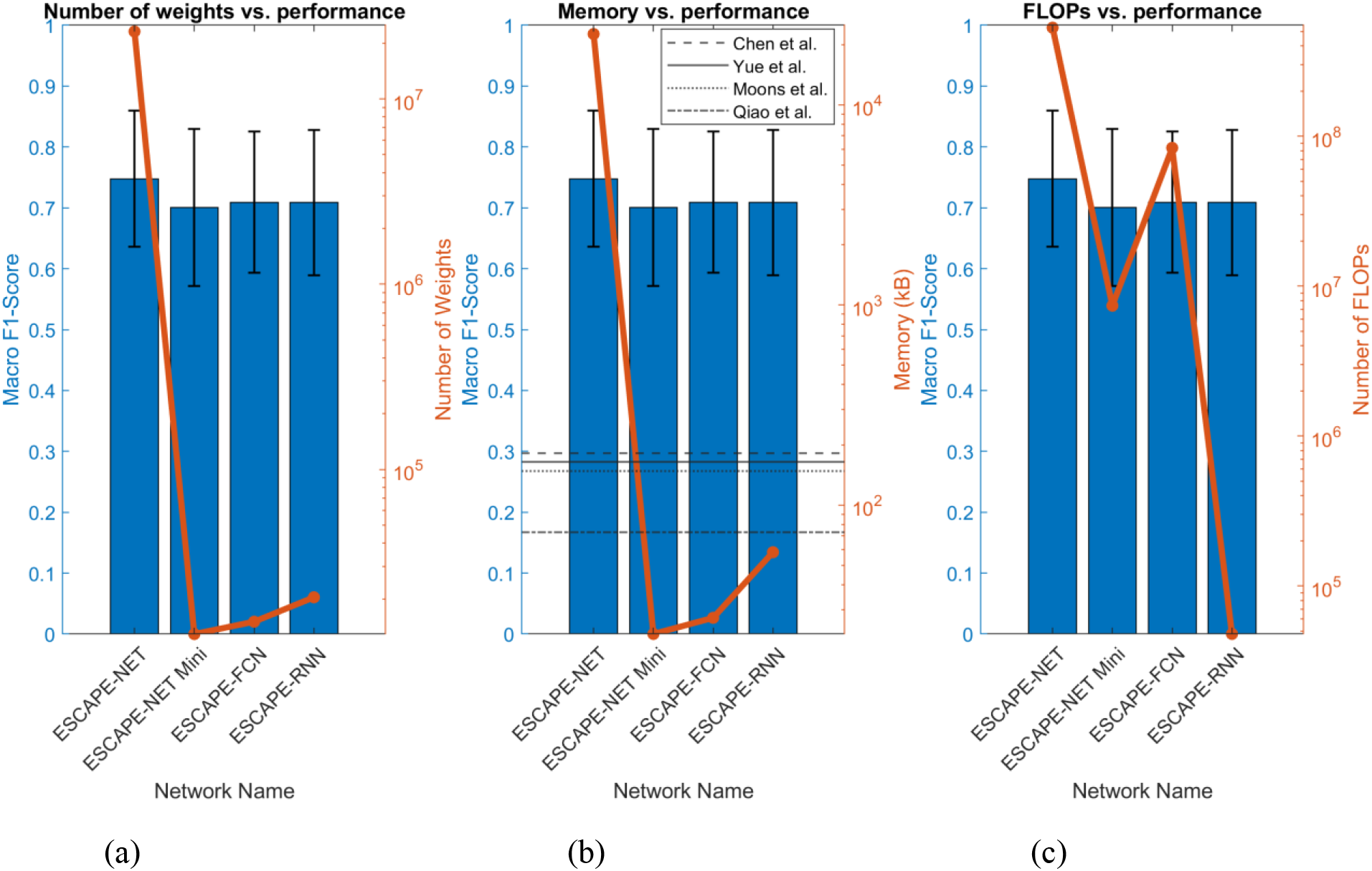
Comparisons between each highlighted network configuration with the baseline ESCAPE-NET network from [14]. (a) Comparison of number of weights and macro F1-score across all 4 networks. (b) Comparison of memory footprint and macro F1-score across all 4 networks. The horizontal dotted lines represent the memory requirements from cited works [30]-[33] respectively. (c) Comparison of number of FLOPs and macro F1-score across all 4 networks.

Power consumption is a related metric to memory use and impacts the usability of implantable devices. Power consumption of neural networks depends on characteristics of the hardware platform on which they are implemented and the energy required to read weights from memory and perform each operation [34]. This study’s focus on decreasing the number of weights and FLOPs will have direct impact on decreasing the power consumption of the networks once implemented on a hardware platform.

### 4.3 Defining performance constraints

Defining a particular performance requirement for this technology in a clinical setting will be highly subject and application based. A macro F1-score threshold of 0.7 was defined for this study based on the baseline macro F1-score of 0.747. This value is based on an average over 9 subjects, but this technology’s intended clinical use will be based on performance for individual subjects. Figure 6 shows the resource/performance trade-off of the explored networks using the best and worst performing subjects rather than by averaging all subjects. The worst performing subjects exhibited far lower macro F1-scores with values nearer to 0.40. For these subjects, a larger network may be required for closed-loop FES to succeed. Conversely, the best performing subjects exhibited higher macro F1-scores with values nearer to 0.90. For these subjects, a smaller network may be sufficient for closed-loop FES to succeed. Given this variance in potential ideal networks for each subject, implementation of the neural network on a wearable device or cloud storage may be more practical due to less rigid constraints on network size.

**Figure 6.**
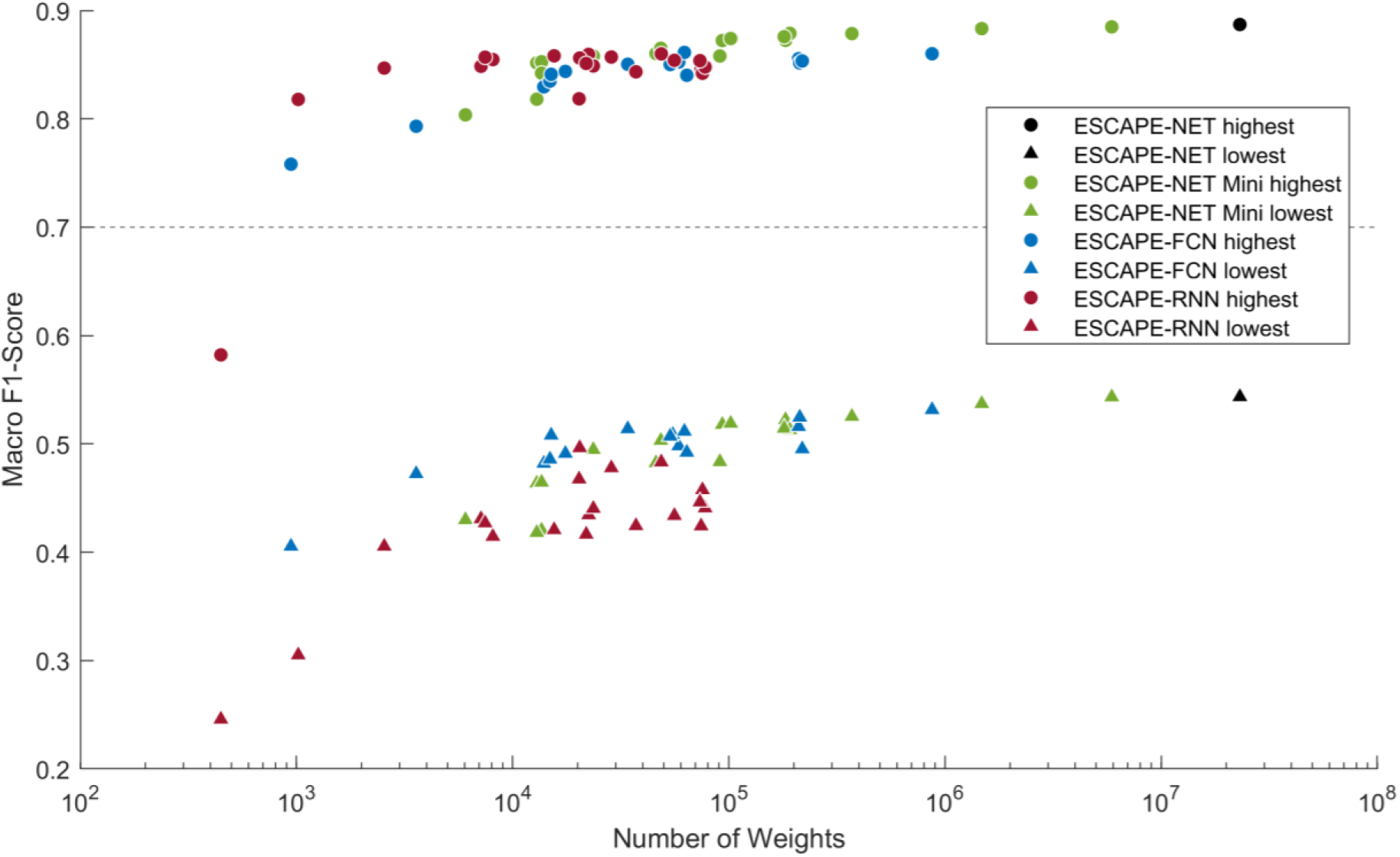
Resource/performance trade-off plot for every explored network configuration for each of the 3 architectures. Each data point represents either the best or worst performing subject out of the 9 subjects used, using the average performance of the networks trained during 3-fold cross-validation. The dotted line represents the performance constraint used in this study.

### 4.4 Network convergence inconsistencies

Due to the stochastic nature of neural network training, sometimes a network failed to converge for certain subjects. The 3 highlighted networks in this study converged for all subjects and folds. In this study, a network’s final performance achieved after either the maximum number of training epochs or the early stopping criterion was met was incorporated for analysis regardless of whether or not the network converged.

### 4.5 Future work

This demonstration used the rat sciatic nerve due to its simplicity for an initial proof of concept. This method should be applied to more complex nerves, such as the human median nerve, to expand the potential reach of such technology. This demonstration was based on acute studies as a proof of concept for the proposed technology. Future work should progress towards chronic studies as this technology is intended for eventual chronic use.

This study has demonstrated that more efficient machine learning techniques are effective at neural pathway discrimination. Future work should focus on implementing these neural networks on ASIC technology that are suitable for surgical implantation. The problem definition addressed in this work was whether NNs with reduced resources could perform with minimal performance loss compared to the original architecture. Future work should focus on performance improvements given resource constraints, potentially using a cost function to ensure an optimal solution is reached.

## 5 Conclusion

This study demonstrates that it is possible to achieve resource-efficient neural networks that perform with minimal accuracy loss compared to ESCAPE-NET and that are suitable for implementation on resource constrained implantable hardware. These efficient networks require a fraction of the resources spent by ESCAPE-NET, requiring 1,132-1,787x fewer weights, 389-995x less memory, and 6-11,073x fewer FLOPs, while maintaining macro F1-scores of over 0.70. The efficient networks have memory requirements within the constraints set by several published deep learning accelerators in 65nm ASIC fabrication technology, which is the targeted platform on which these efficient networks are intended to be implemented.

The ability of efficient neural networks to discriminate between different types of peripheral nerve signals using a single extraneural device is a crucial step towards implementing closed-loop FES technology on implantable devices. Sophisticated closed-loop FES technology can restore nuanced natural control of paralyzed limbs, and implementation on implantable devices can ensure that the developed assistive device is not visible and does not affect the patient’s quality of life. Beyond the impact on people with paralyzed limbs and spinal cord injury, this selective peripheral nerve recording technology has applications in the control of prosthetic limbs and neuromodulation interventions for diseases such as chronic pain, diabetes, or incontinence.

